# Cell fate determination is associated with changes in competing transcriptional units in the human *GATA1* and *GATA2* lineage-determining transcription factors

**DOI:** 10.1101/2024.11.08.622131

**Authors:** Erik M Anderson, Hongchuan Li, Michael Ruesch, Paul W Wright, Aharon G Freud, Stephen K Anderson

## Abstract

The GATA1 and GATA2 transcription factors play a central role in early cell fate decisions in hematopoietic progenitor cells. Although the switch from GATA2 to GATA1 occupancy at GATA-binding sites in erythroblast-specific genes has been extensively studied, the underlying molecular mechanisms controlling this switch are not fully elucidated. An antisense promoter in the 5’ region of the *GATA2* gene produces a long non-coding RNA that has been shown to affect *GATA2* transcription and erythroblast differentiation. The recent identification of an antisense promoter in the first intron of the *GATA1* gene indicates that similar promoter competition mechanisms operate in these genes, potentially controlling GATA1/GATA2 levels in a probabilistic manner. In the current study we perform a comprehensive evaluation of *GATA1* and *GATA2* transcripts in human CD34^+^ progenitors either freshly isolated or differentiated in vitro and the human leukemia cell lines K562 and HL60. The ratio of competing sense and antisense transcripts varied significantly with differentiation status, suggesting that promoter competition controls cell fate decisions. Treatment of HL60 cells with differentiating agents resulted in significant changes in the ratio of *GATA2* sense to antisense promoter activity, in agreement with previous results showing that sense and antisense transcription are associated with distinct cellular phenotypes.

## Introduction

Differentiation of multipotent progenitors is a complex process involving multiple signaling pathways, transcription factors, alternative splicing, post-transcriptional regulation, chromatin-modifying enzymes, DNA methylase/demethylases and additional epigenetic factors. Although the process of cell differentiation has been extensively studied, the mechanism controlling the frequency at which competing cell phenotypes are generated is poorly understood.

Studies of the class I MHC receptors expressed by natural killer (NK) cells have indicated promoter competition as a mechanism controlling variegated gene expression. A bidirectional promoter active in immature NK cells was shown to program the frequency of *Ly49* gene expression in mice (Saleh et al. 2004) and a proximal bidirectional promoter together with an opposing intronic antisense promoter were found to control the human *KIR* gene family encoding MHC receptors (Li et al. 2008; Wright et al. 2013). The relative strength of the competing sense and antisense promoters in *Ly49* and *KIR* genes correlates with the frequency of gene activation, and single nucleotide polymorphisms in transcription factor (TF) binding sites can alter relative promoter strength and change the expression frequency.

A recent investigation of bidirectional promoters in human transcription factors (TFs) revealed that both convergent and divergent antisense transcripts are enriched in TFs, suggesting that promoter competition may allow for variegated expression of TFs (Li et al. 2023).

The mouse and human *GATA2* genes share a similar structure, with a distal hematopoietic/neural cell specific promoter (IS) and a proximal promoter (IG) active in most GATA2-expressing cells (Minegishi et al., 1998; Pan et al., 2000). Examination of the *GATA2* gene in the UCSC Genome Browser (Kent et al. 2002) and the Eukaryotic Promoter Database (EPD) (Dreos et al. 2015) reveals the presence of an additional intermediate promoter in the human *GATA2* gene, however no studies related to this promoter have been published.

GATA2 is a master regulator of hematopoiesis necessary for the maintenance of hematopoietic stem cells (HSCs) and multipotent hematopoietic progenitors. In addition, it is required for the normal development of erythroid precursors, mast cells, megakaryocytes, and eosinophils (Bresnick et al. 2010; Suzuki et al. 2013). Knockout of GATA2 in hematopoietic progenitors greatly reduces the number of HSCs and is sufficient to prevent mast cell differentiation, but terminal megakaryocytic and erythroid differentiation can still occur (Tsai and Orkin 1997). Erythroid and megakaryocyte development is instead controlled by GATA1. GATA1 is found in common myeloid progenitors, eosinophils, mast cells, erythroblasts, and megakaryocytes. Knockout of GATA1 stops erythropoiesis at the proerythroblast stage and results in increased proliferation and decreased maturation of megakaryocyte progenitors (Ferreira et al. 2005; Suzuki et al. 2013). The *GATA1* gene can generate two GATA1 isoforms: either a predominant full-length isoform (GATA1-FL) that initiates translation in exon 2 of the gene; or an alternative isoform that produces a truncated protein called GATA1 short or GATA1s which uses an alternative start site in exon 3 (Shimizu and Yamamoto 2016). GATA1s is missing the first 83 amino acids in its N-terminus and lacks the N-transactivation domain. Although GATA1s possesses similar DNA binding activity as compared to its full-length counterpart, it has greatly reduced transactivation activity (Takasaki and Chou 2024). While a small amount of GATA1s is made in healthy individuals, the expression of GATA1s alone can result in hematological abnormalities and impaired erythropoiesis. *Gata1s* mutant mice only capable of producing the short isoform were found to have a reduced number of erythroid cells and an increased megakaryocyte population with no significant change in their common progenitor population as evaluated by differential gene expression analysis. *Gata1s* mice also had dysregulated global H3K27 methylation and impaired terminal differentiation of erythroid cells (Ling et al. 2019). In humans, exclusive production of GATA1s causes similar impairments in erythropoiesis and is seen in diseases such as GATA1 related Diamond-Blackfan anemia and myeloid leukemia of Down syndrome (Gialesaki et al. 2018; Takasaki and Chou 2024).

During normal hematopoiesis, GATA1 and GATA2 need to be temporally regulated. GATA2 is required early and GATA1 is present later as cells adopt erythroid or megakaryocytic cell fates. GATA2 activates both the *GATA1* and *GATA2* genes while GATA1 activates *GATA1* but inhibits *GATA2* expression resulting in a dynamic change in gene expression called GATA switching (Grass et al. 2003; Bresnick et al. 2010). The full-length GATA1 protein is required for GATA switching, as GATA1s mice have impaired switching and higher GATA2 levels (Ling et al. 2019). GATA1 can repress *GATA2* by interacting with FOG1 (Friend of GATA-1; *ZFPM1* gene). FOG1 is a coregulator of GATA proteins and can interact with the N-finger domains of both GATA1 and GATA2. FOG1 is required to facilitate chromatin occupancy of GATA1 and displace GATA2 at the *GATA2* locus to repress its activity as GATA1 mutants unable to interact with FOG1 are incapable of repressing *GATA2* during erythroid commitment (Crispino et al. 1999; Bresnick et al. 2010; Chlon and Crispino 2012). Interestingly, FOG1 is sufficient to prevent mast cell differentiation, as overexpression of FOG1 in mast cell progenitors redirected them into erythroid, megakaryocytic, and granulocytic cell lineages (Cantor et al. 2008).

While the mechanism of GATA switching and the role of FOG1 in hematopoiesis have been extensively studied, relatively little research has been done on the regulatory role of the antisense transcripts of the *GATA1* and *GATA2* genes. The *GATA2-AS1* transcript is a long noncoding RNA (lncRNA) with a promoter that opposes the distal IS promoter of the *GATA2* gene (Li et al. 2023). A recent study has implicated the *GATA2-AS1* transcript in the regulation of erythroblast differentiation. Deletion of the first exon of this transcript in HUDEP2 erythroid progenitor cells caused a loss of detectable antisense, and resulted in both accelerated erythroid differentiation and dysregulation of erythroblast gene expression (Liu et al. 2024). Additional work on *GATA2-AS1* in colorectal cancer cell lines showed that the *GATA2* antisense was able to interact with DDX3X to stabilize *GATA2* mRNA which in turn upregulated *GATA2-AS1* (Pan et al., 2022). A patient with a duplication of the entire *GATA*2 locus leading to *GATA2-AS1* overexpression presented a classical GATA2 deficient phenotype (Singh et al. 2021). Lastly, differential gene expression analysis of 5’scRNA-seq data from cultured CD34^+^ cells comparing cells that expressed *GATA2* sense only vs cells containing *GATA2* antisense only, showed that *GATA2* sense cells had enrichment of mast/basophil related genes while *GATA2* antisense only cells had enrichment of erythroid genes (Li et al. 2023). These data were also analyzed to compare *GATA1* sense only cells against *GATA1* antisense only cells and found that *GATA1* sense only cells expressed erythroid genes as expected. However, *GATA1* antisense only cells displayed a mast/basophil phenotype and had increased *GATA2* sense expression (Li et al. 2023). *GATA1* antisense contains two exons that overlap with exons in the upstream suppresser of variegation 3-9 homolog 1 (*SUV39H1)* gene, a histone methyltransferase that trimethylates lysine 9 of histone 3 (H3K9) and is responsible for gene silencing and heterochromatin formation. H3K9 methylation is known to increase significantly as cells differentiate and is a major epigenetic hurdle for *in vitro* induction of pluripotency (Onder et al. 2012; Padeken et al. 2022). The mouse *Suv39h1* gene was shown to be an indirect target of the pluripotency TF OCT4, which activates the promoter of a *Suv39h1* antisense lncRNA containing the same two antisense exons as the *GATA1* antisense transcript, repressing *Suv39h1* sense expression and maintaining cells in an undifferentiated state (Bernard et al. 2022). *GATA1* antisense may play a similar role in hematopoiesis, as K562 cells with a deletion in the *GATA1* antisense promoter had increased *SUV39H1* transcription, indicating a similar inhibitory role for *GATA1* antisense, necessitating additional research (Li et al. 2023).

The current study provides a detailed evaluation of the transcriptional landscape of the human *GATA1* and *GATA2* genes using 5’-directed single cell RNA sequencing of ex vivo isolated CD34^+^ progenitor cells, in vitro differentiated CD34^+^ cells, the human K562 erythroleukemia cell line and the HL60 myeloid leukemia line. The relative levels of sense and antisense transcription varied among different cell types. Differentiation of the HL60 cell line into monocytes was associated with increases in the relative level of *GATA2* sense transcription, whereas differentiation into granulocytes was associated with increased antisense transcription, implicating promoter competition in cell fate determination.

## Results

### Evaluation of sense and antisense promoter/enhancer elements in the GATA1 gene

The regulation of the *GATA1* gene during hematopoiesis has been extensively studied in mice, revealing multiple cis-acting elements (Vyas et al. 1999). Some of the mouse elements have been shown to be conserved in the human gene, such as the HS1 (−3.5) and HS3 (+3.5) elements (Valverde-Garduno et al. 2004). However, the recent discovery of a strong antisense promoter in the first intron of the *GATA1* gene has indicated that additional studies of the human *GATA1* transcriptional landscape are needed (Li et al. 2023). 5’-directed single cell RNA sequencing of freshly isolated human CD34^+^ cells, differentiating CD34^+^ cultures, and the HL60 myeloid and K562 erythroid leukemia cell lines was performed to assess *GATA1* gene transcription at different stages of differentiation. Figure 1 shows read density tracks taken from an Integrative Genome Viewer (IGV) (Robinson et al. 2011) analysis of single cell libraries. Human elements with homology to the mouse HS1 and HS3 regions were found to produce spliced *GATA1* transcripts, raising the possibility that these elements may function as promoters and/or enhancers depending on the state of cell differentiation. Transcripts originating from upstream elements amounted to less than one percent of total *GATA1* transcription and were only observed in cell populations with high levels of transcripts from the predominant *GATA1* promoter (Pro1). Sense transcripts from the downstream elements were rare, despite the DE1 element constituting a bidirectional element with strong antisense transcription. Interestingly, transcripts from Pro1 and the upstream elements spliced into either the 2^nd^ or 3^rd^ exon at differing ratios, while transcripts from the downstream elements only spliced into the 2^nd^ exon. This indicates that the relative levels of GATA1-FL and GATA1s are not fixed and may change during cell differentiation.

**Figure 1.**
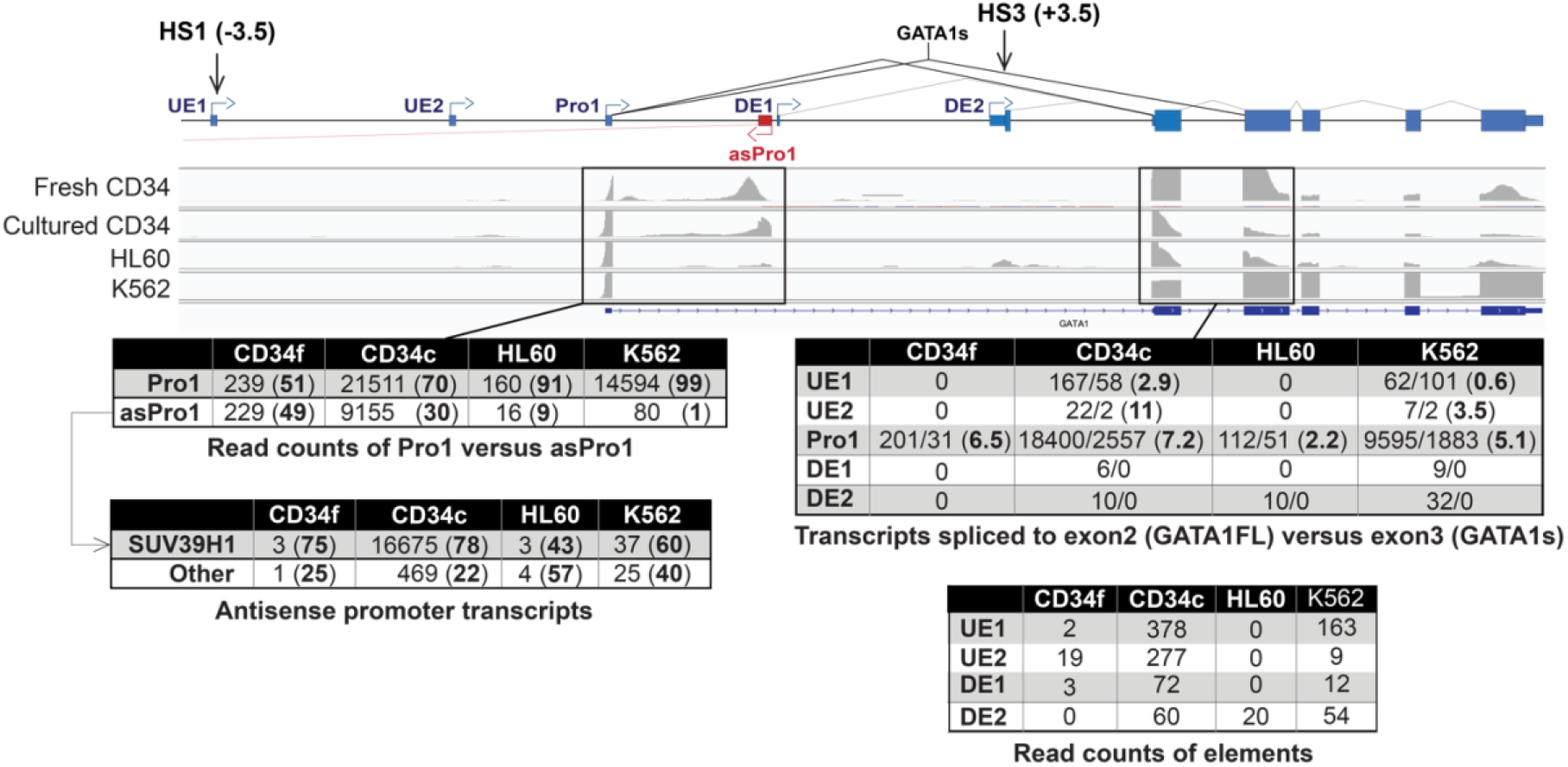
The ratio of *GATA1* sense to antisense is highly variable. IGV coverage tracks from single cell 5’-RNAseq of freshly isolated CD34^+^ cells (Fresh CD34), a 2-week culture of CD34^+^ cells (Cultured CD34), the HL60 cell line, and the K562 cell line are shown for the *GATA1* gene. A schematic of the promoter/enhancer elements identified is shown *above* the IGV tracks. Upstream elements (UE) and downstream elements (DE) relative to the *GATA1* promoter (Pro1) are shown. The previously characterized HS1 (−3.5) and HS3 (+3.5) elements are indicated. The read counts of the competing Pro1 and Pro1as promoters are boxed and enumerated in the table on the *left below*. The observed read counts of each promoter are shown, with the percentage of read counts for each promoter indicated in bold in parentheses. An additional table evaluating the splicing of antisense transcripts is shown *below*, and the percentage of transcripts splicing into the *SUV39H1* gene or other exons is indicated in bold in parentheses. The boxed region on the *right* and accompanying table shows the read counts of transcripts spliced to the 2^nd^ or 3^rd^ exon, and the ratio of transcripts splicing to the 2^nd^ exon, which are capable of producing the full-length protein versus transcripts that splice to the 3^rd^ exon and code for GATA1s are shown in bold in parentheses. The additional table below evaluates the read counts observed for the UE and DE elements.

The relative activity of the *GATA1* Pro1 and opposing intron 1 antisense promoter (asPro1) varies significantly between cell types, with fresh CD34^+^ cells having nearly equivalent activity, differentiating CD34^+^ cultures shifting toward sense transcription, and the leukemic cell lines showing dominant sense activity. The majority of antisense transcripts splice into the *SUV39H1* gene, potentially helping to maintain an undifferentiated state. However, most of the asPro1 transcripts in fresh CD34^+^ cells initiated downstream of the splice donor site and were not spliced, producing a relatively short lncRNA. This suggests that antisense transcripts in the fresh CD34^+^ cells are able to inhibit *GATA1* transcription without affecting SUV39H1 expression, potentially enabling differentiation into non-erythroid cell types.

### Evaluation of sense and antisense promoter elements in the GATA2 gene

Although three sense promoters and one antisense promoter have been identified in the human *GATA2* gene, 5’ RNAseq analysis of CD34^+^ cells, differentiating CD34^+^ cell cultures, as well as the HL60 and K562 cell lines revealed the presence of additional spliced antisense transcripts originating from the intermediate (Pro2) and proximal IG (Pro3) promoters, indicating their bidirectional nature (Figure 2). All three antisense transcripts share an identical 2^nd^ exon, however there are three potential 3^rd^ exons that are used to varying extents by the antisense transcripts. The level of the novel antisense transcripts varied between cell types, ranging from 5 to 20% of the previously characterized *GATA2-AS1* transcript. Interestingly, there was significant variation in the levels of antisense transcripts and the distal *GATA2* Pro1 (IS) transcripts in different cell types, indicating cell type specificity and potential competition between these opposing elements. The levels of competing *GATA2* Pro1 and asPro1 transcripts varied between cell types, as seen for *GATA1* sense/antisense promoter activity. However, the observed patterns were quite different: the *GATA2* sense transcripts were dominant in the CD34^+^ cells, sense and antisense were balanced in HL60 cells, and there was a striking shift toward antisense in K562 cells.

**Figure 2.**
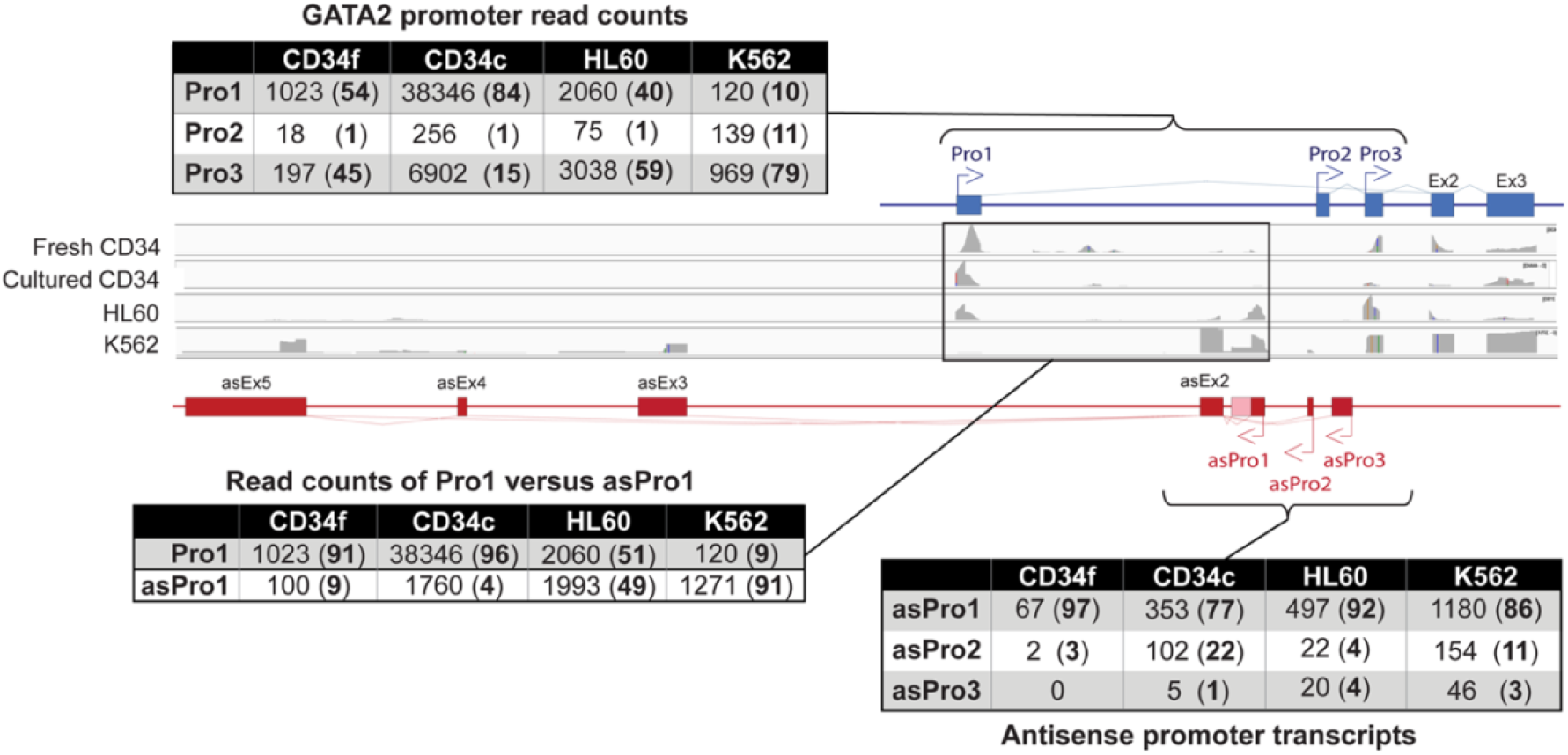
*GATA2* sense versus antisense activity changes with cell differentiation status. IGV coverage tracks from single cell 5’-RNAseq of freshly isolated CD34^+^ cells (Fresh CD34), a 2-week culture of CD34^+^ cells (Cultured CD34), the HL60 cell line, and the K562 cell line are shown for the *GATA2* gene. A schematic of *GATA2* sense promoter elements identified is shown *above* the IGV tracks, and antisense elements are shown *below*. The relative activity of each sense promoter based on read counts in the first exon is shown in the table *above*, with the percentage of transcripts indicated in parentheses in bold. The boxed region and corresponding table *below* on the *left* evaluates the relative activity of the Pro1 sense versus the asPro1 promoter by comparing read counts of the first exon. Percentages are shown in bold in parentheses. A table evaluating the number of spliced antisense transcripts from each antisense promoter is shown *below* on the *right*, with percentages shown in bold in parentheses.

### In vitro analysis of GATA2 promoter activity

In vitro promoter assays were performed to confirm the bidirectional promoter activity associated with the novel antisense transcripts identified and determine if the relative sense versus antisense activity varies between different cell lines as observed for the bidirectional promoters identified in the *AHR, GATA3*, and *RORC* genes (Li et al. 2023). Figure 3 compares sense and antisense promoter activity of luciferase reporter constructs containing fragments encompassing the bidirectional promoter region or the upstream IS promoter (Pro1). The distal Pro1 promoter exhibited minimal antisense promoter activity, consistent with the lack of detectable antisense transcripts from this promoter region. In contrast, the proximal Pro3 (IG) promoter showed varying levels of antisense promoter activity, with high antisense activity correlating with high sense activity in the different cell lines. Although sense transcription was dominant, the ratio of sense to antisense varied among the cell lines, suggesting that differences in transcription factor expression might influence the sense to antisense ratio.

**Figure 3.**
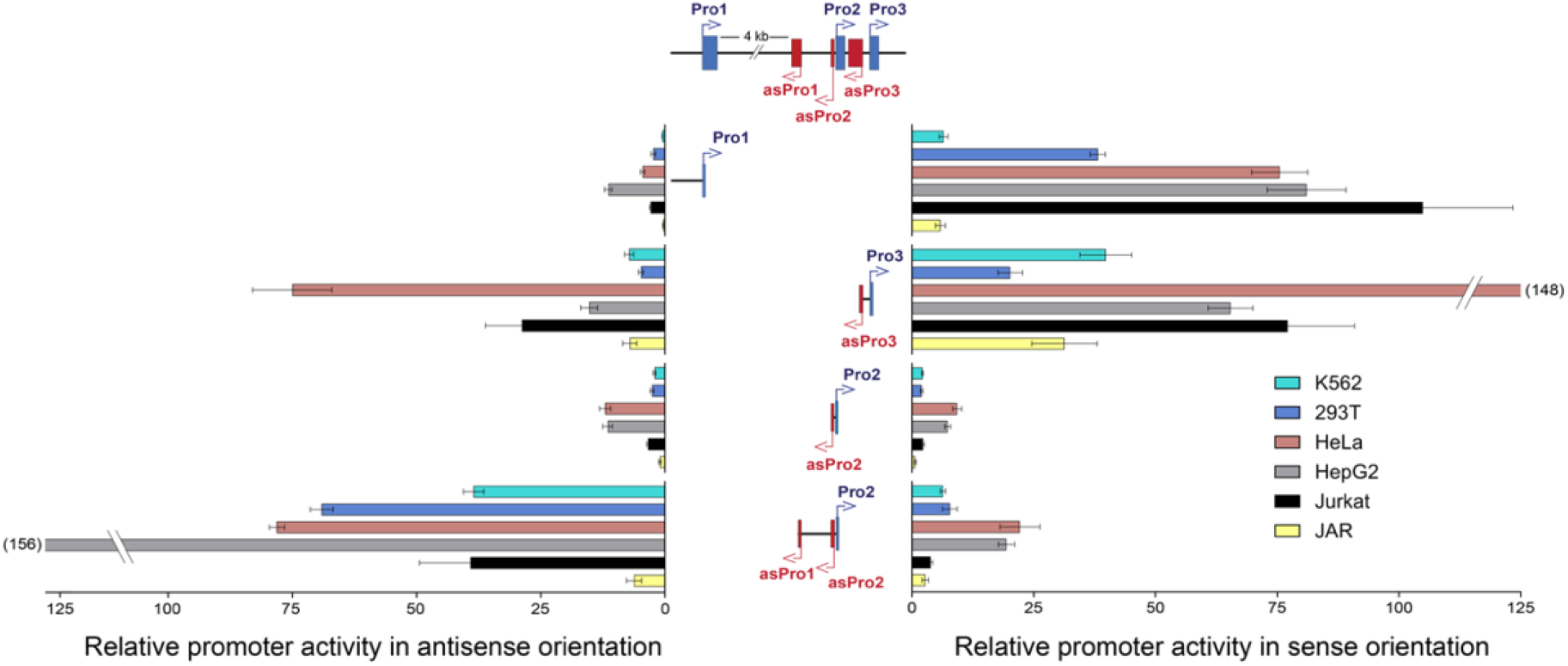
*In vitro* promoter activity of *GATA2* luciferase constructs. Luciferase promoter assay of eight pGL3 constructs evaluating the sense (*right graph*) and antisense (*left graph*) activity of four regions in the *GATA2* gene across six cell lines representing multiple tissue types. The x-axis for both graphs depicts the fold change in light units relative to the empty pGL3 vector for their respective directions, and the promoter pair is shown on the y-axis. *Above* the two graphs is a simplified model of the *GATA2* gene which displays the relative positions of the sense promoters (blue arrows) and the antisense promoters (red arrows) as well as the first exon of each promoter. The *central* schematic indicates the location of each pGL3 construct, as well as the promoters included as indicated by the attached arrows. Values represent the mean and standard error of the mean of 3 to 5 independent experiments.

A total of six cell lines were analyzed representing multiple tissue types, including bone marrow (K562), embryonic kidney (HEK293T), uterine (HeLa), liver (HepG2), peripheral blood T lymphoblast (Jurkat), and placenta (JAR). HL60, a promyeloblast cell line, was not included since it was not amenable to luciferase experimentation due to high background luminescence. The ratio of sense to antisense activity at each of the promoter regions tested was consistent between samples as seen in Table 1, with the small bidirectional Pro3/asPro3 and Pro2/asPro2 promoter pairs particularly consistent. However, there were a few key differences between cell lines when comparing which sense or antisense promoter was dominant. When analyzing the ratio between Pro1 and Pro3, it was found that HEK293T and Jurkat cells preferred Pro1, mirroring what was seen in the scRNAseq data in cultured CD34^+^ cells (Figure 2). K562, HeLa, HepG2, and JAR primarily used Pro3, mirroring the fresh CD34^+^ and HL60 RNAseq data. Antisense Pro1 (asPro1) was the dominant antisense promoter in K562, HEK293T and HepG2 cells, as it was in the RNAseq data of all 4 cell types analyzed; the ratio of asPro1 to asPro3 was about equal in HeLa, Jurkat, and JAR cells (Figure 3).

**Table 1.**
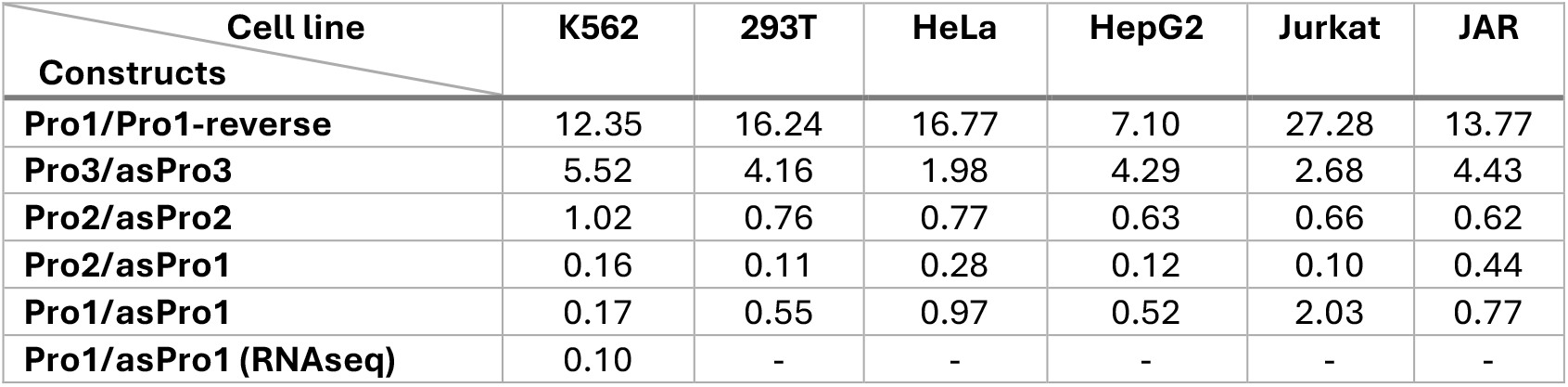
The ratio of sense to antisense luciferase activity by cell type, related to figure 3. The first four rows show the ratio of sense activity versus antisense activity for each of the four bidirectional promoter elements within the *GATA2* region. The fifth row instead compares the convergent pair Pro1 versus asPro1. The sixth row shows the ratio of *GATA2* Pro1 reads to asPro1 reads observed in the K562 RNAseq data (see Figure 2).

| Cell line<br>Constructs | K562 | 293T | HeLa | HepG2 | Jurkat | JAR |
| --- | --- | --- | --- | --- | --- | --- |
| Pro1/Pro1-reverse | 12.35 | 16.24 | 16.77 | 7.10 | 27.28 | 13.77 |
| Pro3/asPro3 | 5.52 | 4.16 | 1.98 | 4.29 | 2.68 | 4.43 |
| Pro2/asPro2 | 1.02 | 0.76 | 0.77 | 0.63 | 0.66 | 0.62 |
| Pro2/asPro1 | 0.16 | 0.11 | 0.28 | 0.12 | 0.10 | 0.44 |
| Pro1/asPro1 | 0.17 | 0.55 | 0.97 | 0.52 | 2.03 | 0.77 |
| Pro1/asPro1 (RNAseq) | 0.10 | - | - | - | - | - |

The most notable difference between all cell types tested was the ratio of transcription of the competing convergent promoters Pro1 and asPro1. Considering the read counts of Pro1 versus asPro1 observed in Figure 2, sense transcription was shown to be dominant in both fresh and cultured CD34^+^ cells which is also seen to a lesser extent in the Jurkat luciferase data. HeLa cells were found to have relatively similar sense and antisense activity, as seen in the HL60 RNAseq data, whereas asPro1 was more active than Pro1 in HEK293T and HepG2 cells. Notably, K562 was distinct, as it had dominant antisense activity, with approximately sixfold higher antisense activity observed in the luciferase data and a tenfold difference when comparing read counts from Figure 2. Collectively, these data indicate significant changes in the activity of sense relative to antisense *GATA2* transcription across different tissue types.

### In vitro differentiation of the HL60 cell line correlates with changes in relative Pro1 sense versus Pro1 antisense transcription in the GATA2 gene

Previous results have implicated competition of sense and antisense transcription in the *GATA1* and *GATA2* genes in the generation of distinct cell phenotypes in differentiating CD34^+^ hematopoietic progenitor cells. To determine if similar mechanisms could operate in the in vitro differentiation of a leukemic cell line, we used 5’-directed single cell sequencing to look for changes in TF transcription in HL60 cells induced to differentiate into either monocytes or granulocytes. Figure 4A shows the single cell RNAseq results obtained after overnight treatment of HL60 cells with either DMSO or PMA to induce granulocyte or monocyte differentiation respectively. The UMAP depicts the separate clustering of control HL60 cells versus the DMSO or PMA treated groups. RT-PCR with primers for the granulocyte-specific genes *ARG1* and *CEACAM8*, or the monocyte-specific *DCSTAMP* gene showed expression in the appropriate treatment group (Supplemental Fig. S3), and mapping of the transcripts onto the UMAP identified a subpopulation of cells expressing the genes in each cluster. Differential gene expression analysis revealed that both *GATA2* sense and antisense transcripts were significantly upregulated in PMA-treated cells, warranting further examination. Analysis of *GATA2* transcription in differentiating HL60 cells using IGV revealed significant differences in the relative levels of *GATA2-AS1* transcripts as compared to Pro1 or Pro3 sense transcripts (Figure 4B). When comparing total sense versus antisense transcription, control HL60 cells were sense dominant, whereas DMSO-treated cells were antisense dominant. PMA-treated cells were strongly sense dominant with sense transcripts far exceeding the level of antisense transcripts. These differences were even more apparent when comparing the ratio of Pro1 to asPro1 across the three treatment groups. Control cells expressed both transcripts almost equally with a ratio of 1.0, DMSO-treated cells had over eightfold more asPro1 transcripts compared to Pro1, while PMA-treated cells had almost three times as many Pro1 transcripts. These results suggest that high *GATA2* sense transcription drives monocyte differentiation, whereas high antisense transcription favors granulocyte production.

**Figure 4.**
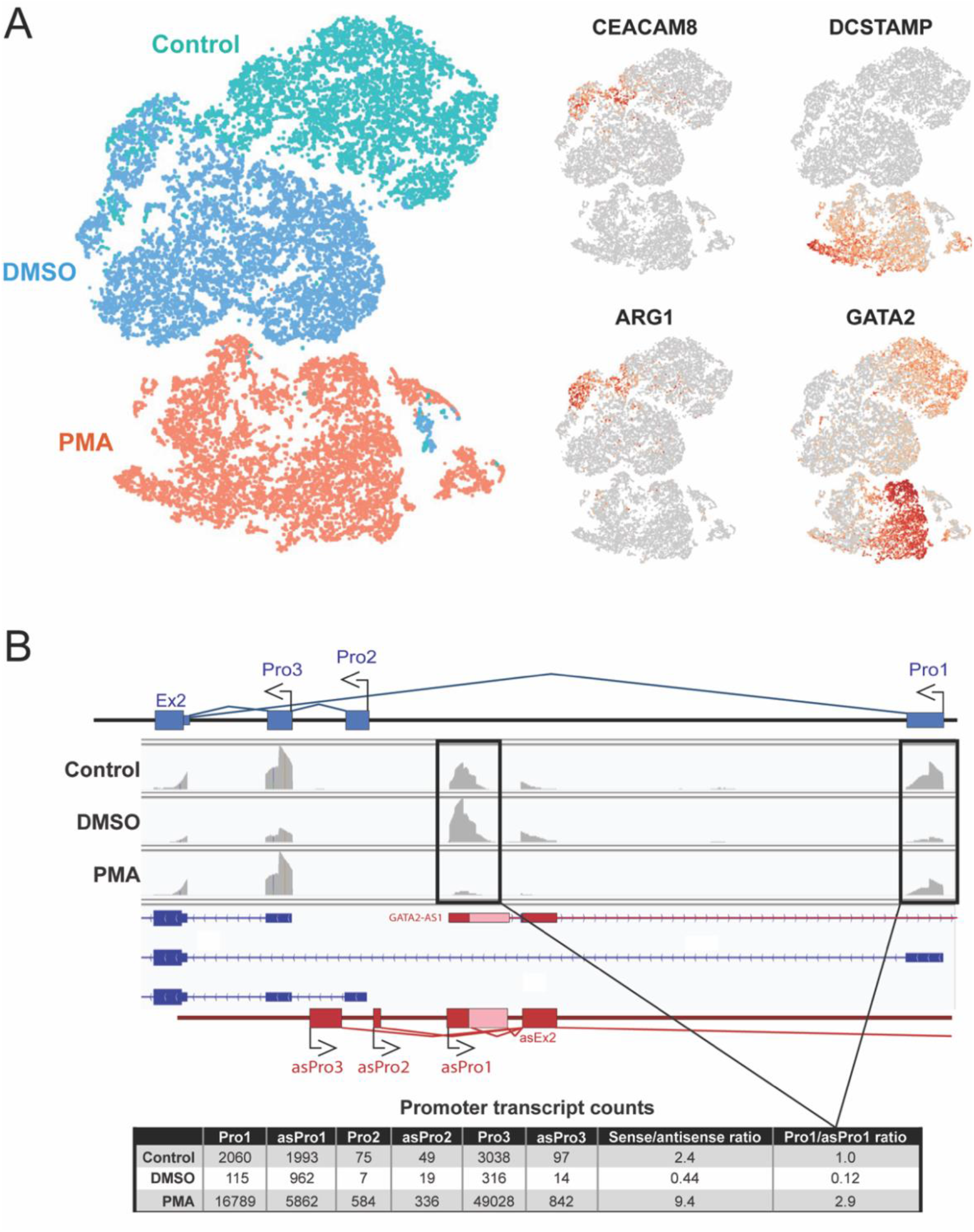
In vitro differentiation of HL60 cells changes the ratio of sense to antisense transcription in the *GATA2* gene. (*A)* Aggregated UMAP visualized in the Loupe Browser (10x Genomics) of single cell 5’-RNAseq data of HL60 cells. The larger (*left*) schematic separates the cells by treatment group. Untreated control cells are in green, DMSO-treated cells are in blue, and PMA-treated cells are in orange. The four smaller schematics (*right*) show the relative expression of granulocyte markers *CEACAM8* and *ARG1*, monocyte marker *DCSTAMP*, and *GATA2*. (*B*) IGV coverage tracks from single cell 5’-RNAseq of the *GATA2* promoter region in HL60 cells for each of the three treatment groups. The location of the sense promoters is shown *above* the tracks in blue and the antisense promoters are shown *below* in red. The lines between exons indicate splicing patterns. The light pink rectangle indicates alternative splicing of the asPro1 exon. The Pro1 and asPro1 reads are boxed, and the table *below* shows the transcript counts for each promoter, the ratio of total sense to antisense counts, and the ratio of Pro1 to asPro1 counts in each treatment group.

## Discussion

Detailed analyses of the *GATA1* and *GATA2* genes and the role of cis-acting elements in hematopoiesis has identified multiple enhancer regions that are important for normal cell differentiation. The promoter region and upstream *GATA1* HS1 element are sufficient for expression in primitive erythroid cells, while the *GATA1* HS3 element in the first intron is required for efficient expression in definitive erythroid cells (Onodera et al. 1997). These elements are considered to act as tissue-specific enhancers. However, the observation of full-length spliced transcripts from these elements in the current study indicates promoter activity. In fact, since the K562 single-cell library was sequenced using PacBio technology, the results reveal that 161 of the 164 observed transcripts originating from the HS1 (UE1) element in K562 cells represent full-length spliced cDNAs, clearly indicating promoter function (Figure S1). Promoters and enhancers share many common attributes, and some strong promoters have enhancer activity (Andersson and Sandelin 2020). Although enhancers produce short bidirectional RNAs (eRNAs), one of the distinguishing features of promoters is the production of long spliced RNAs. The production of full-length transcripts from the *GATA1* elements would therefore classify them as promoters. However, the level of transcription from the HS1 and HS3 elements represents less than 2% of the level of transcription from the principal gene promoter, and thus would have little impact on protein expression directly, but may impact chromatin structure. The conservation of these elements between mouse and human indicates an important function that may require the generation of full-length spliced transcripts. Perhaps transcription from these elements plays a role in the initial activation of the gene. Alternatively, low level transcription may enhance the specificity of gene expression by disrupting downstream promoter complexes, requiring their reassembly, similar to the kinetic proofreading mechanism proposed to increase specificity of enhancers and promoters (Boeger 2022). In contrast to the weak promoter activity of the UE1, UE2, and DE2 elements, the DE1 (asPro1) intron 1 antisense promoter activity is comparable to sense Pro1 promoter activity, and it produces spliced transcripts capable of silencing the *SUV39H1* gene, clearly indicating that it functions primarily as a promoter element (Li et al. 2023). Sense transcription from this element can produce spliced full-length transcripts, however, they are extremely rare, constituting less than 0.1% of the antisense transcripts from asPro1, indicating that this is primarily a unidirectional antisense promoter. In freshly isolated CD34^+^ cells, most asPro1 transcripts are unspliced, terminating near the first exon and thus would be expected to compete with the *GATA1* promoter without inhibiting *SUV39H1*. Since *SUV39H1* inhibition has been associated with maintenance of the progenitor state (Onder et al. 2012; Bernard et al. 2022), cells producing unspliced asPro1 transcripts may be directed into non-erythroid cell lineages due to inhibition of GATA1 expression. The cultured CD34^+^ cells produced primarily spliced antisense transcripts, the majority of which included two exons antisense to *SUV39H1* exons (Figure 1; Genbank# PQ241452). The *GATA2* gene has two well-characterized enhancer elements located at -3.9 kb (Sanalkumar et al. 2014) and +9.5 kb (Johnson et al. 2012; Hsu et al. 2013; Soukup and Bresnick 2020) relative to the major transcription start site of the gene. The -3.9 element confers chromatin accessibility at this site but is dispensable for *Gata2* expression. The +9.5 element was found to play an essential role in embryogenesis and hematopoiesis. In contrast to the enhancer elements described in the *GATA1* gene, no spliced transcripts were observed to originate in the *GATA2* enhancer elements. There were however, short enhancer-associated transcripts detected, consistent with the identification of these elements as enhancers (Supplemental Fig. S2).

The detection and characterization of elements associated with open chromatin likely depends on the differentiation status of the cells that are analyzed. For example, a distal *Ly49* gene element (−4.5 kb) was originally characterized as an enhancer required for expression in NK cells (Tanamachi et al. 2004). However, studies of NK progenitors and an immature NK cell line revealed that this element functions as a promoter producing full-length *Ly49* transcripts in immature NK cells (Saleh et al. 2002, 2004). A similar observation was made for the *GATA1* HS1 (UE1) element in the current study. Short transcripts flanking the element were observed in fresh CD34^+^ cells as would be expected for an enhancer element, whereas the majority of transcripts observed in differentiating CD34^+^ cells and the K562 line were spliced (Supplemental Fig. S1). Therefore, one would consider this element as an enhancer in the CD34^+^ progenitors, but as a promoter in the differentiating erythroblasts or the K562 erythroleukemia cell line.

The functions of GATA1 and GATA2 and their interplay during hematopoiesis has been extensively studied in many biological systems. In comparison, the role of *GATA* antisense transcripts in hematopoiesis has received relatively little attention. However, in recent years there has been accumulating evidence supporting functional roles for these antisense RNAs. For example, GATA3 acts as a master regulator of T-cell development that downregulates the Th1 (T helper type 1) pathways and is necessary for Th2 terminal differentiation (Scheinman and Avni 2009). The divergent *GATA3* antisense transcript (*GATA3-AS1*) is elevated in Th2 cells and was found to be required for the normal expression of *GATA3*, augmenting its expression (Gibbons et al. 2018). The knockdown of *GATA6-AS1*, the divergent antisense transcript of *GATA6*, in embryonic stem cells effectively blocked endoderm differentiation due to decreased GATA6 levels, showing that *GATA6-AS1* is necessary for normal GATA6 expression (Yang et al. 2020). A recent study by Liu et al knocked out *GATA2-AS1* in HUDEP-2 cells, an erythroid progenitor cell line, and found the knockout reduced GATA2 expression, and KO cells differentiated more rapidly and were less proliferative compared to wildtype controls (Liu et al. 2024). All these studies describe similar mechanisms regarding the relationship between the sense and antisense transcripts of *GATA* genes, and conclude that they mostly work cooperatively to promote differentiation.

We took an agnostic approach to the HL60 differentiation model, unbiased as to what our genes of interest would be, looking instead for which genes had switching activity between sense and antisense when adopting different cell fates, which led to the identification of the *GATA2* sense/antisense switch. In our proposed model, the competition between *GATA2* sense and antisense acts as a binary switch controlling cell fate. Sense dominant HL60 cells become monocytes, antisense dominant cells become granulocytes, and undecided cells express both sense and antisense about equally. However, moving forward it will be important to knock out *GATA2* asPro1 in HL60 cells to verify if removing antisense transcription changes how they differentiate when treated with PMA or DMSO. If this model is correct, it is likely the HL60 cells lacking the asPro1 promoter will become resistant to granulocyte differentiation and become predisposed to adopt monocyte cell fates. In addition, it will be of interest to perform knockouts of intronic antisense promoters like *GATA2 asPro1* in lineage determining TFs in CD34^+^ progenitor cells to verify if these antisense promoters affect lineage decisions in ex vivo-derived human cells. It has been established that more than 25% of human and mouse genes contain antisense transcripts overlapping sense exons in coding genes (Anderson and Anderson 2024). Therefore, it is possible that the novel sense/antisense switch described here could be active in a variety of biological processes worth further examination.

## Materials and methods

### Cell lines and Cell culture

HeLa and HEK293T cells were cultured in Dulbecco’s modified Eagle’s medium and HepG2 cells were cultured in Eagle’s minimum essential medium. The Jurkat and JAR cell lines were maintained in RPMI1640 Medium. HL60 and K562 cells were cultured in Iscove’s Modified Dulbecco’s Medium (IMDM; ATCC). Each culture medium contained 10% fetal bovine serum, 100 U/ml each of penicillin and streptomycin, sodium pyruvate, and L-glutamine. All cell lines were grown in cell culture incubators at 37°C in the presence of 5% CO2.

### CD34^+^ cell isolation and culture

Apheresis cones were flushed using 10 mL of PBS +1% FBS + 2mM EDTA and incubated on a nutator at room temp for 20 min with 500 μL of RosetteSep Human NK Cell Enrichment Cocktail (Stem Cell Technologies). Following the incubation, the cells were diluted in PBS and layered over Ficoll-Paque PLUS (GE Healthcare) and centrifuged at 2000 rpm for 20 min with the brake off. Enriched mononuclear cells were then collected, and residual RBCs were lysed using eBioscience 1X RBC lysis buffer (ThermoFisher Scientific). RBC lysis buffer was diluted in PBS, and cells were re-pelleted at 1800 rpm for 5 min. CD34^+^ hematopoietic progenitor cells were enriched using the Indirect Human CD34 MicroBead Kit (Miltenyi Biotec). Following cell enrichment, cells were stained for flow cytometry and CD34^+^ cells were further purified via sorting to >99.9% purity on an FACSAria II cell sorter (BD Biosciences). The culture media for CD34^+^ cell support following isolation contained DMEM and F12 (2:1 ratio) supplemented with 1% antibiotic/antimycotic (ThermoFisher Scientific), 20 mg/mL ascorbic acid, 24 μM 2-ME, 0.05 mg/mL sodium selenite (Sigma-Aldrich), and 10% heat-inactivated human AB serum (Valley Biomedical). CD34^+^ cells were plated in media supplemented with 10 ng/mL each of human Flt3 ligand, c-Kit ligand, and IL-3 (Peprotech).

### Generation of luciferase reporter plasmids

Promoter fragments from the *GATA2* gene were generated by PCR with ZymoTaq PCR Master Mix (Zymo Research) using the primers listed in Table 2. The PCR products were ligated using GenBuilder Cloning Kit (Genscript) into linearized pGL3-Basic (Promega) vector cut with XhoI/HindIII (NEB) to generate constructs in both forward and reverse orientations. Each construct was verified by Sanger sequencing. Plasmid DNA for each verified construct was isolated using ZymoPURE II Plasmid Midiprep kit (Zymo Research).

**Table 2.**
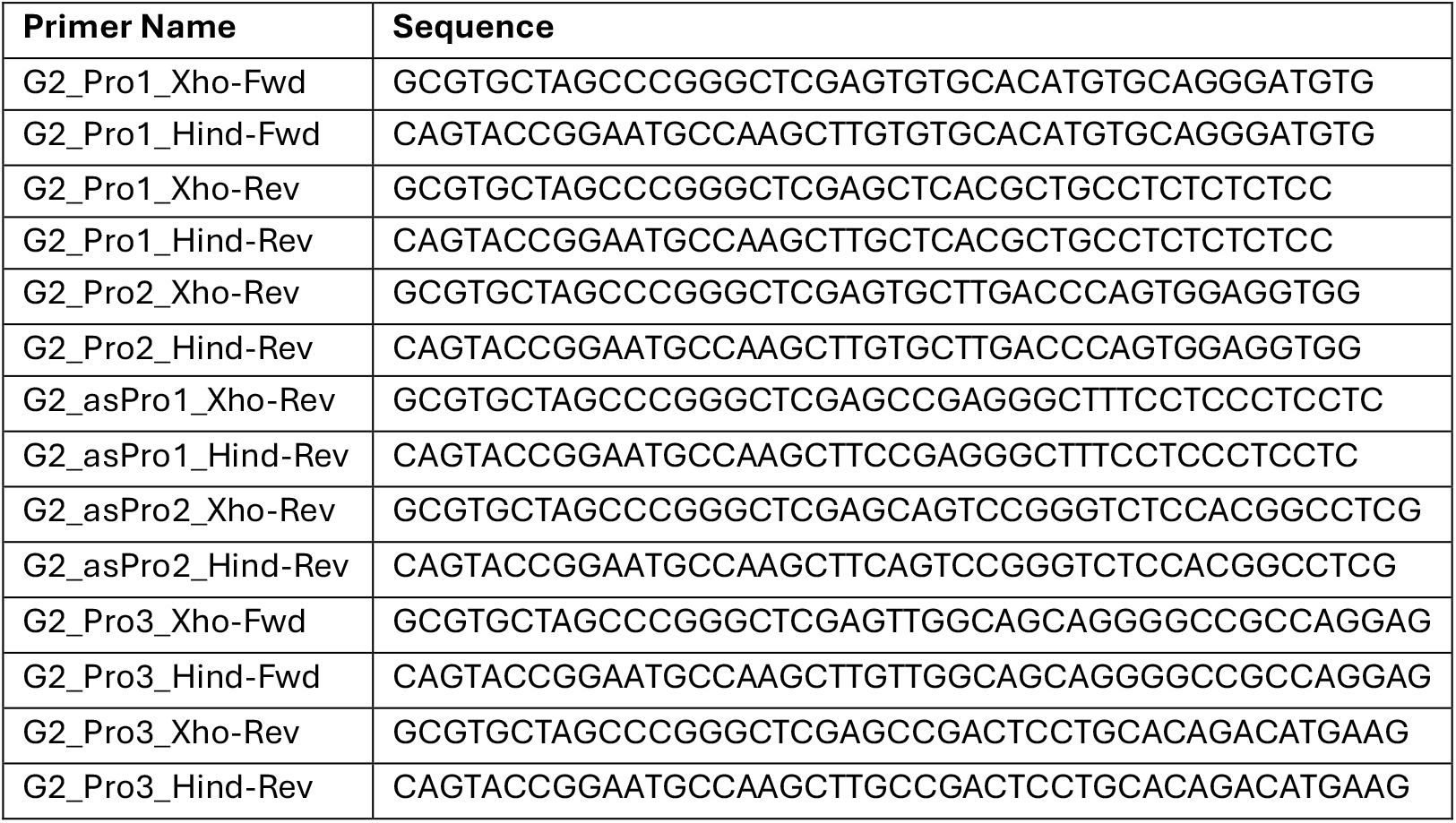
Primers used to amplify *GATA2* promoter fragments for cloning into pGL3.

### Cell transfection and luciferase reporter assay

HEK293T cells were plated at 1.0×10^5^ cells per well in a 24-well plate the day before transfection and incubated overnight at 37°C in 5% CO_2_. For each well, 200 ng of pGL3 reporter construct plus 10 ng of Renilla luciferase pRL-SV40 control DNA was diluted in 50 μL of buffer. 1 μL of jetOPTIMUS (Genesee Scientific) transfection reagent was added and incubated at room temperature for 10 min. The DNA mixture was then added to each well and incubated at 37°C in 5% CO_2_ for 48 h before analysis. HeLa and HepG2 cells followed the same general protocol but instead were plated at 2.5×10^5^ cells per well in a 24 well-plate, and 1 μg of of pGL3 reporter construct, 50 ng of Renilla and 2 μL of jetOPTIMUS (Genesee Scientific) transfection reagent was added.

For suspension cell line transfection, 5×10^5^ cells per well were seeded for both Jurkat and JAR cells in a 24-well plate the day before transfection. For both cell lines, 2 μg of reporter construct DNA, 200 ng of Renilla DNA, 4 μL of jetOPTIMUS transfection reagent were diluted in 100 μL buffer and incubated at room temperature for 10 min before addition to each well. For K562 cells, 2×10^5^ cells per well were plated in each well of a 24-well plate the day before transfection. For each well, 2 μg of pGL3 reporter construct and 100 ng of Renilla was added to 50 μL of OPTI-MEM Reduced Serum medium (Gibco) and 1.25 μL of Plus Reagent (Invitrogen) and incubated at room temperature for 15 minutes. 2.5 μL of LTX Reagent was then added (Invitrogen) and the mixture was incubated for an additional 25 minutes before being added to each well. Each treated well was then and incubated at 37°C in 5% CO_2_ for 48 h before analysis.

Luciferase activity was assayed using the Dual-Luciferase Reporter Assay System (Promega) according to the manufacturer’s instructions. Briefly, the culture medium was removed 48 h post-transfection and cells were washed with 0.5 mL of phosphate buffered saline (PBS, pH 7.4). Cells were then lysed with passive lysis buffer. The suspensions were centrifuged at 10000g for 1 min. A total of 20 μL of cell lysate supernatant was mixed with 100 μL of luciferase substrate, and the light units were measured on a luminometer (Promega). Measurements of the firefly luciferase activity of the promoter constructs were normalized relative to the activity of the Renilla luciferase produced by the pRLSV40 control vector and each construct was tested in duplicate in at least three independent experiments.

### HL60 Differentiation

HL60 cells were plated at 4.0×10^5^ cells per well in a 6-well plate the day before treatment and incubated overnight in 4 mL of IMDM medium at 37°C in 5% CO_2_. The following morning each well was collected, the media was removed and the cells were washed with 1 mL of phosphate buffered saline (PBS, pH 7.4). The control cells were then replated in 4 mL of fresh media. For the DMSO treated cells, 100 μL of Dimethyl Sulfoxide Cell Culture Reagent (MP Biomedicals) was added to 4 mLs of media comprising 2.5% of the total volume. For PMA treated cells, 8 μL of 10 μg/mL PMA (InvivoGen) dissolved in DMSO was added to 4 mLs of media for a final concentration of 20 ng/mL or approximately 32 nM PMA. All three treatment groups were incubated 24 h at 37°C in 5% CO_2_. The following morning the control and DMSO treated cells were pelleted by centrifugation and resuspended in 1 mL media for transport. Since the PMA treated cells became adherent, they were first trypsinized for 5 minutes at 37°C in 5% CO_2_. The PMA treated cells then were pelleted and resuspended in 1 mL of untreated media. The three treatment groups were then subjected to single cell RNA sequencing.

### Single Cell RNA sequencing

HL60 and K562 cells were washed twice with PBS + 0.04% BSA at room temperature, resuspended in 500-1000 μl of the same buffer and counted. The cell viability of the samples was 93%-96%. Approximately,10,000 HL60 cells were loaded on the 10X Genomics Chromium platform to target ∼10,000 cells and libraries were constructed with Chromium Next GEM Single Cell 5’ Reagent Kits v2 (Dual Index) according to the manufacturer’s instructions. The cDNA quality and quantity were determined using Agilent Tapestation 4150 (Agilent Technologies Inc). The HL60 libraries were sequenced on NovaSeq S1 100 cycles asymmetric paired-end run with read length of 10 bp for the sample index reads and 150 bp for the Read 1 and Read 2 to achieve a sequencing saturation about 70%. The freshly isolated CD34^+^ cells were cultured overnight, centrifuged at 300g for 5 minutes, resuspended in PBS + 1% FBS and 2mM EDTA for staining with TotalSeq™-C antibodies (Biolegend). Following staining, the cells were washed 3 more times and 2,500 cells from each of four donors were loaded on the 10X Genomics Chromium platform and libraries were constructed with a Chromium Next GEM Single Cell 5′ reagent kit v2 dual index according to the manufacturer’s instructions. For K562 cells, approximately 15,000 cells were loaded on the 10X Genomics Chromium platform to target ∼10,000 cells using Chromium Next GEM Single Cell 5’ Reagent Kits v2 (Dual Index). The single-cell MAS-Seq libraries were constructed with the Kinnex single-cell RNA kit (Pacific Biosciences) according to the manufacturer’s instructions. The sequencing primer was annealed and Revio polymerase was bound to the library before loading using the Revio Polymerase kit (Pacific Biosciences). The libraries were sequenced on Revio SMRTcell at 150 pM loading concentration. Sequencing was performed during a 24-hour movie time on Revio.

### PacBio MAS Single Cell Analysis

PacBio HiFi reads were processed using the PacBio SMRT link analysis pipeline (smrtlink-release_13.1.0.221970). The HiFi reads were segmented into individual segmented reads (S-reads) that represent the original cDNA sequences using Skera (1.2.0). Primers and polyA tails were removed using lima (2.10.0). Iso-seq pipeline was run to extract single-cell barcode and UMI information. The cell barcodes were corrected according the 10X genomics single cell whitelist. Cell barcodes that represent encapsulated single cells (as opposed to ambient RNA) were also identified using the percentile method. Reads are then deduplicated based on cell barcodes and UMIs. Deduplicated reads were mapped using phmm2 (1.14.0) to the reference genome (refdata-gex-GRCH38-2020-A) and classified against a transcript annotation (GENCODE version 44). For transcript classification and annotation, pigeon was used (1.2.0) which is based on SQANTI3 to annotate the identified splice variants for transcripts in each cell. Gene- and isoform-level single-cell matrix files were output for tertiary analysis. In addition, Seurat (version 5.0.1) and singleR (2.4.1) were run to perform gene count matrix clustering and cell type annotation.

### Evaluation of relative promoter activity in single cell 5’-RNAseq data

The capture of the 5’ ends of transcripts in the single cell RNAseq libraries examined allows for a comparison of relative promoter activities by evaluating relative read counts in the first exon associated with each promoter. The coverage track in IGV was used to determine the level of transcription associated with the first exon of each promoter. The maximum read counts observed in a 50-nucleotide window adjacent to the principal transcription start site was recorded for each promoter, and the relative percentage was calculated.

## Data availability

Single cell RNA sequencing data for cultured CD34^+^ cells are available at the NCBI Gene Expression Omnibus under accession number GSE242100. Additional data or reagents are available from the corresponding author upon request.

## Competing interest statement

The authors claim no competing interests

## Acknowledgments

This project has been funded in whole or in part with federal funds from the Frederick National Laboratory for Cancer Research, National Institutes of Health, under contract HHSN261200800001E. This research was supported in part by the Intramural Research Program of NIH, Frederick National Lab, Center for Cancer Research. Funding was also provided from extramural NIH/NCI grant CA208353 to A.G.F. We would like to thank the CCR Sequencing Facility at the Frederick National Laboratory for Cancer Research for performing single-cell capture and sequencing.

## Author Contributions

S.A. and A.F. conceived the study. E.A., H.L., M.R, and P.W. generated the data. S.A. and E.A. analyzed the data. S.A., and E.A. wrote the manuscript. S.A. and A.F. acquired the funding.

## Notes

### Competing Interest Statement

The authors have declared no competing interest.

