## Supplemental Fig1 for "Cell fate determination is associated with changes in competing transcriptional units in the human *GATA1* and *GATA2* lineage-determining transcription factors"

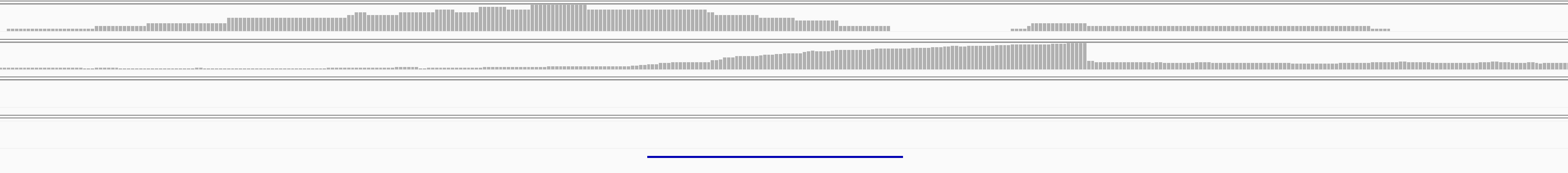

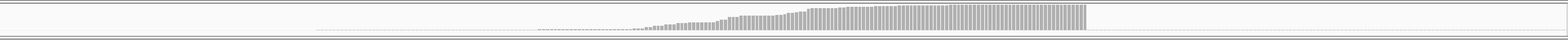

Fresh CD34

Cultured CD34

HL60

K562

UE1 exon

**Figure S1.** (related to Figure 1). The *GATA1* UE1 element functions as a promoter in GATA1-expressing cells. Coverage tracks of genomic regions flanking previously characterized *GATA1* -3.5 kb UE1 element are shown. The location of the UE1 exon identified is shown below.
