## Supplemental Fig2 for "Cell fate determination is associated with changes in competing transcriptional units in the human *GATA1* and *GATA2* lineage-determining transcription factors"

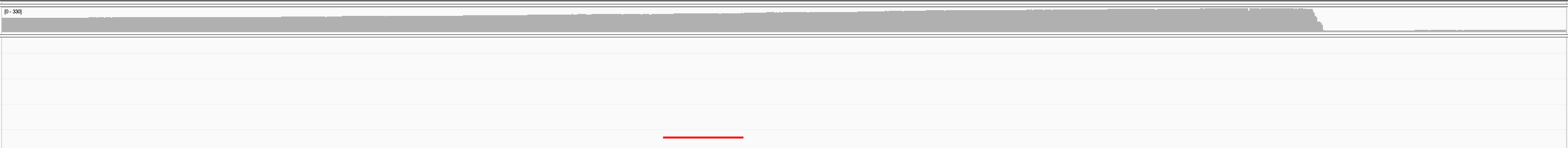

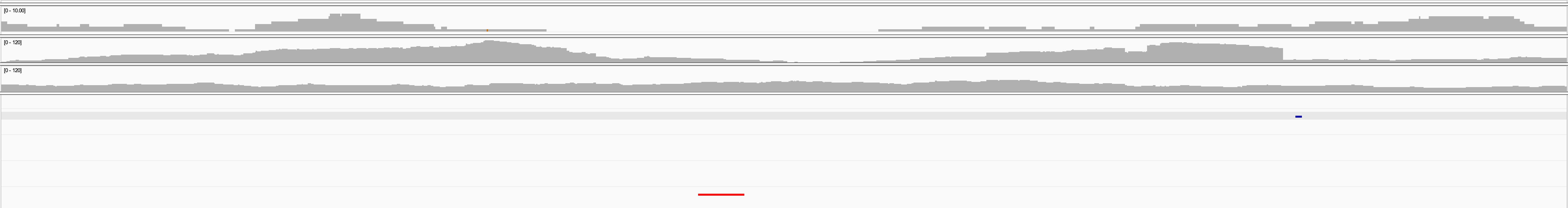

-3.9

Enhancer

1 kb

Fresh CD34

Cultured CD34

HL60

K562

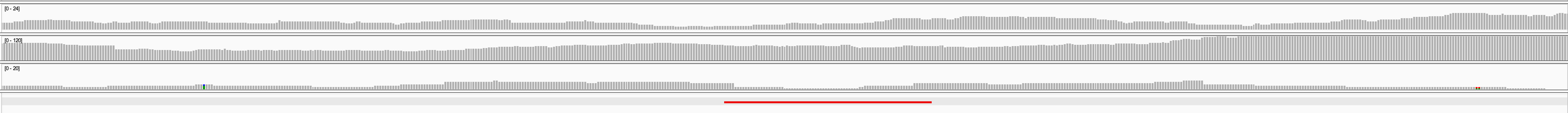

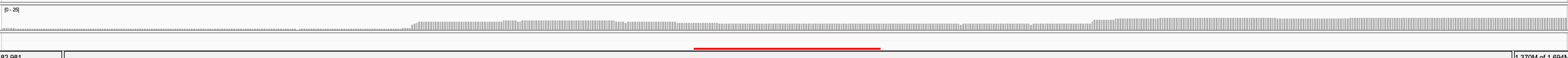

+9.5

Enhancer

500 bp

A

B

Fresh CD34

Cultured CD34

HL60

K562

**Figure S2**. (related to Figure 2). Detection of short enhancer-associated RNAs in *GATA2.* Coverage tracks of genomic regions flanking previously characterized *GATA2* -3.9 kb (*A*) and +9.5 kb (*B*) enhancer elements are shown. The location of human sequences homologous to the mouse enhancers is indicated by the black bars beneath each panel.
