## Supplemental Fig3 for "Cell fate determination is associated with changes in competing transcriptional units in the human *GATA1* and *GATA2* lineage-determining transcription factors"

**No RT Control**

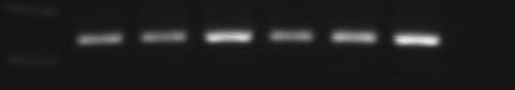

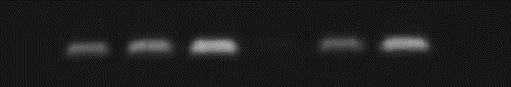

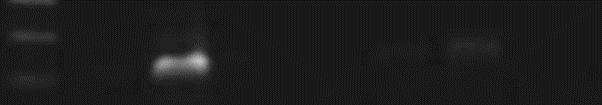

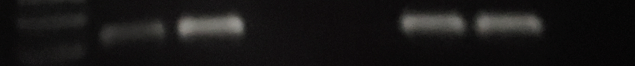

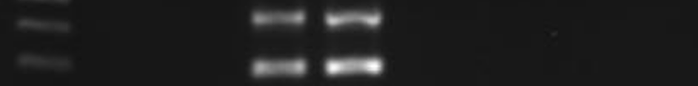

**GATA2**

**GATA2-AS**

**ARG1**

**CEACAM8**

**DCSTAMP**

**DMSO 24 hours**

**DMSO 72 hours**

**PMA 24 hours**

**PMA 72 hours**

**Control 24 hours**

**Control 72 hours**

**Figure S3.** (related to Figure 4). RT-PCR analysis of genes associated with HL60 differentiation. Results of an RT-PCR analysis of genes associated with granulocyte (*ARG1*, *CEACAM8*) differentiation after treatment with DMSO, or genes associated with monocyte differentiation (*GATA2*, *DCSTAMP*) after treatment with PMA.
